# First worldwide detection of *bla*_IMP-15_ in *Stenotrophomonas maltophilia* isolated from a patient in Lebanon

**DOI:** 10.1101/2025.02.28.640784

**Authors:** Ziad C. Jabbour, Jose-Rita J. Gerges, Hadi M. Hussein, Sara B. Barada, Lama Hamadeh, Rami Mahfouz, Zeina A. Kanafani, Ghassan M. Matar, Antoine G. Abou Fayad

**Author notes:** Corresponding author (A.G. Abou Fayad).

## Abstract

*Stenotrophomonas maltophilia* is an intrinsically multi-drug resistant (MDR) bacterium initially found in the environment that is emerging worldwide. The rate of isolation from immunocompromised patients combined with limited treatment options make *S. maltophilia* a new concern in clinical settings. Here we report the first detection of the Metallo-β-Lactamase (MBL) gene bla_IMP-15_ in *S. maltophilia* isolated from a patient in Lebanon. The isolate exhibited an extensively drug-resistant profile with high resistance to trimethoprim-sulfamethoxazole, levofloxacin, and minocycline considered by the *Clinical and Laboratory Standards Institute* (CLSI) as first-line antimicrobials used to treat such infections. Resistance to newly adopted anti-*S. maltophilia* agents was noted such as ceftazidime-avibactam. However, the isolate only showed susceptibility to cefiderocol and synergy was observed upon treatment with a combination of ceftazidime-avibactam and aztreonam by disk diffusion. Long-read and short-read whole-genome sequencing was performed and generated a hybrid assembly of 8 contigs. The isolate belonged to strain T50-20 and showed a novel sequence type. Moreover, several antimicrobial resistance genes conferring resistance to multiple antimicrobial classes were found. Particularly, bla_IMP-15_ was detected on insertion sequence IS6100 surrounded by transposition elements. Furthermore, the presence of the IMP gene was confirmed by polymerase chain reaction (PCR) followed by agarose gel electrophoresis and Sanger sequencing. This study highlights that the threat behind the bla_IMP-15_ gene is not only linked to the high resistance to Ceftazidime-avibactam in the *S. maltophilia* isolate but is also of concern due to its transmissibility to other pathogens, conferring alone an MDR profile.

## Importance

Carbapenemases are enzymes that confer resistance to a broad range of antimicrobials, from β-lactams, β-lactams/β-lactamase combinations, cephalosporins, to carbapenems. Recently, the spread of carbapenem resistance has been of great concern according to the World Health Organization (WHO), since carbapenem-resistant organisms are among the critical and high priority bacterial pathogens. Here we report for the first time the presence of the MBL bla_IMP-15_ in *Stenotrophomonas maltophilia* isolated from a patient in Lebanon. *S. maltophilia* is a ubiquitous bacterium that is widely associated with pulmonary diseases especially in the setting of polymicrobial infections. The presence of this bacterium limits treatment options due to its natural resistance to several antimicrobial classes. Studies on this MDR pathogen are scarce and mechanisms of resistance need to be further investigated. Our findings raise concerns about the presence of bla_IMP-15_ in *S. maltophilia* and its potential for dissemination to other organisms thereby spreading carbapenem-resistance in clinical settings.

## Observation

*Stenotrophomonas maltophilia* is a Gram-negative environmental bacterium originating from plant rhizosphere where it thrives as a dominant microorganism in its ecosystem (1). However, in healthcare settings, the isolation of this bacterium is becoming more prominent with cases of respiratory infections and particularly those associated with cystic fibrosis (2). *S. maltophilia* is a globally emerging multi-drug-resistant pathogen. The rise in prevalence of hospital-acquired and community-associated *S. maltophilia* infections is of concern to the public health due to limited treatment options (3). It is well established that this bacterium exhibits intrinsic resistance to several antimicrobial classes including aminoglycosides, cephalosporins, macrolides, tetracyclines, phenicols and carbapenems (4). As such, common treatment options are limited to trimethoprim-sulfamethoxazole, levofloxacin, and tetracycline derivatives. Novel antimicrobial agents such as cefiderocol, newer tetracycline derivatives and novel β-lactam/β-lactamase inhibitor combinations are becoming more clinically useful for treating *S. maltophilia* infections (5). Interestingly, *S. maltophilia* is commonly detected in polymicrobial infections with *Pseudomonas aeruginosa, Klebsiella pneumoniae* and *Acinetobacter baumannii* exacerbating the difficulty in treatment (6,7). As such, the ubiquitous presence of *S. maltophilia* in diverse environments is linked to its ability to acquire genetic material through horizontal gene transfer. Comparative genomic analyses of *S. maltophilia* isolated from various sources concluded that this process plays a critical role not only in its adaptability in different environments, but in the evolution of the dissemination of antimicrobial resistance (8). Particularly, the spread of carbapenem-resistance through carbapenemase-encoding genes, such as IMP-type carbapenemases poses a significant public health threat, limiting treatment options and increasing mortality rates (9). In this paper, we report the first detection of the gene bla_IMP-15_ in *Stenotrophomonas maltophilia* isolated from a tertiary healthcare center in Lebanon.

The *S. maltophilia* isolate STM_202406_001 was recovered from a sputum sample taken from a 67-year-old patient with acute myeloid leukemia initially admitted for chemotherapy and who remained hospitalized due to several complications, namely polymicrobial pneumonia including *Stenotrophomonas* pneumonia, carbapenem-resistant *Enterobacterales* bacteremia, septic shock and *Clostridium difficile* colitis. Antimicrobial susceptibility testing was performed using the Kirby-Bauer disk diffusion method against 12 antimicrobials and 1 combination according to the CLSI guidelines (**Table 1**) (9, 10). The *S. maltophilia* isolate (STM001) showed resistance to antimicrobials previously used and tested: ampicillin, erythromycin, cefotaxime, aztreonam, imipenem, gentamicin, ciprofloxacin according to CLSI guidelines in 2020 (10), and antimicrobials mainly used for treating *S. maltophilia* infections trimethoprim-sulfamethoxazole, levofloxacin, minocycline according to newer guidelines (11) and ceftazidime-avibactam a potent combination and new solution to *Stenotrophomonas maltophilia* infections (12). This extensively drug-resistant isolate showed susceptibility to cefiderocol, a novel siderophore cephalosporin conjugated with a catechol moiety, and an effective antimicrobial agent against IMP-producing *Enterobacterales* (13). In addition, a synergy was observed between aztreonam + ceftazidime-avibactam against this isolate (**Table 1**). To elucidate the molecular basis of this isolate’s resistance profile, whole-genome sequencing was performed.

**Table 1.**
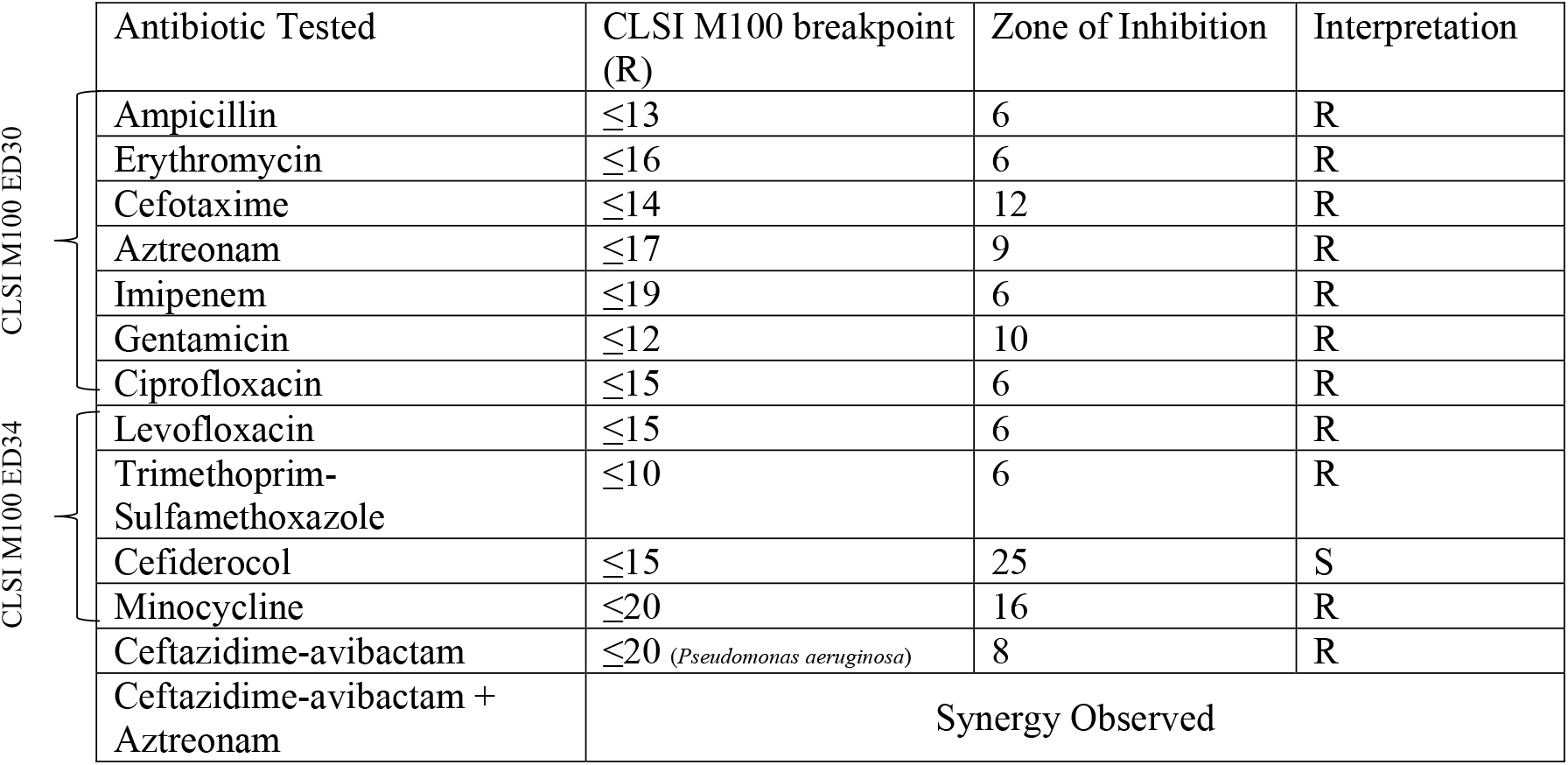
Antimicrobial Susceptibility testing of STM001.

STM001 was subjected to short-read sequencing on the Illumina MiSeq platform (Illumina Inc., San Diego, CA, USA) and long-read sequencing on Oxford Nanopore Technologies MinIon MK1B (Oxford Nanopore Technologies, Oxford, UK). Hybrid assembly of short and long reads was done using the nf-core/bacass (v2.4.0) pipeline (14). Genome annotation was done using and Prokka (v1.14.6) (15). The 741-bp-long metallo-beta-lactamase IMP-15 was identified with 100% identity using CARD (v 3.2.8) (10.1093/nar/gkac920) (16), and NCBI Antimicrobial Resistance Gene Finder (17). The raw sequences were uploaded to NCBI under accession number SAMN46897142.

The whole-genome sequencing revealed 8 contigs with a total of 4,921,506 bp with 4,450 protein-coding genes, 7 rRNA and 72 tRNA with 66% G+C content. STM001 was identified as *Stenotrophomonas maltophilia* strain T50-20 known to have antibacterial properties (18). MultiLocus Sequence Typing (MLST) on STM001 revealed an unknown sequence type with novel alleles of the genes *gapA, guA, mutM* (19). Various antimicrobial resistance determinants were identified including *bla*_IMP-15_ and *bla*_L1_ conferring resistance to carbapenems, *aph(6)-Smalt* and *aph(3’)-IIc* and *aac(6’)-lb-cr* to aminoglycosides, *sul1* to sulfonamides, and *adeF* efflux pump conferring resistance to fluoroquinolones and tetracyclines. The *bla*_IMP-15_ gene was detected downstream of insertion sequence IS6100 along with aac(6’)-lb-cr, *sul1, qacE*, and *aac(6’)-lb3* identified by MGE finder (20). Various transposition tools were surrounding the mobile genetic element that are suspected to have helped with the transposition, these include: Tn3 family transposases ISPa382 and ISPa383, as well as many hypothetical proteins that are potentially involved (Figure 1). No plasmids were noted in the genome.

**Figure 1.**
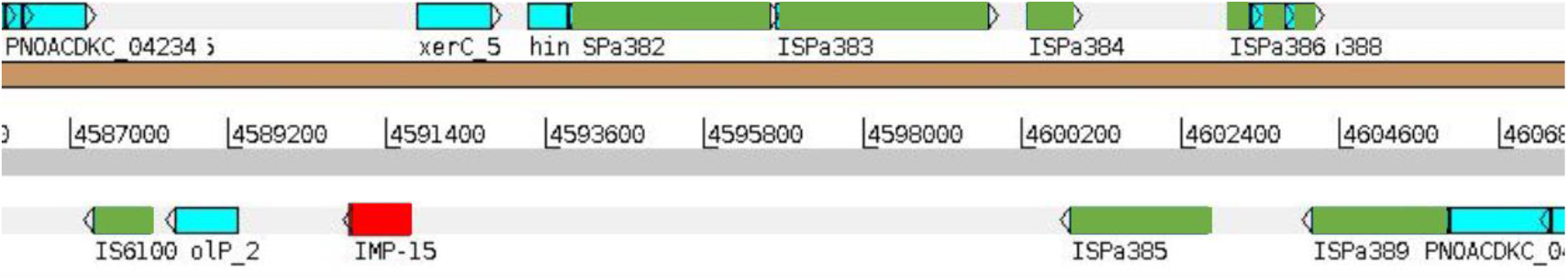
Visualization of the position of the bla_IMP-15_ gene in STM001 on the reverse strand flanked by transposition elements. IS6100 is an insertion sequence encoding a transposase upstream of the gene. ISPa382/383 are transposase-encoding genes, and ISPa385→ISPa389 are hypothetical proteins that could be involved in the transposition.

**Figure 2.**
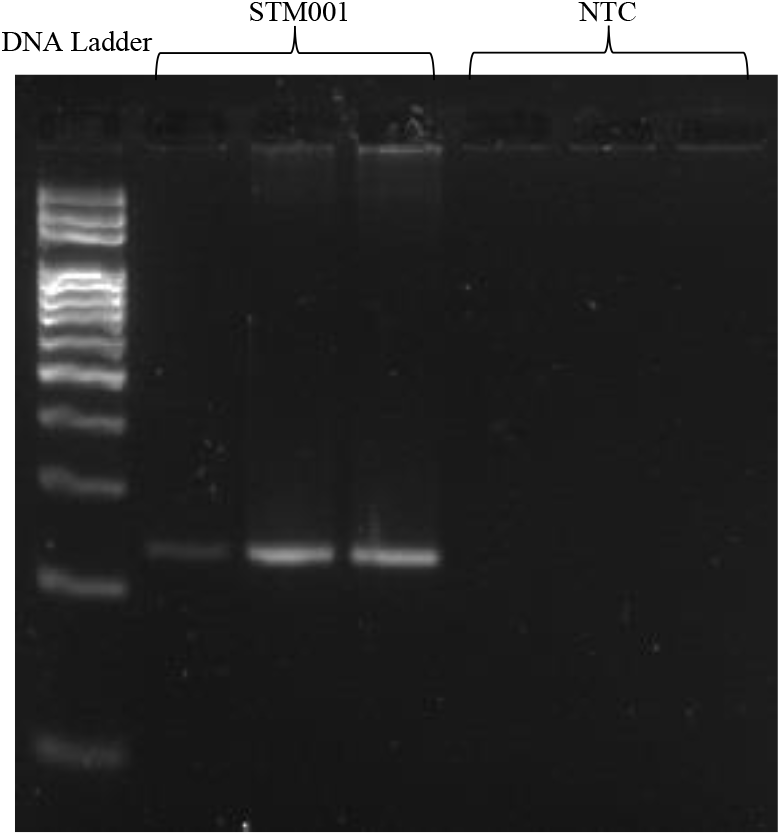
Agarose gel electrophoresis of the bla_IMP-15_ gene purified PCR product. The 1^st^ lane shows the DNA ladder separation, 2^nd^, 3^rd^ and 4th lanes show the purified and amplified IMP-15 gene fragments. Whereas the 5^th^, 6th and 7^th^ lanes show the absence of products in the non-template control replicates.

For further confirmation, a conventional IMP-targeted PCR was performed using the primers: IMP-Forward: *GGAATAGAGTGGCTTAATTCTCTC*

IMP-Reverse: *GGTTTAACAAAACAACCACC*

The same sample was run in triplicate using Solis Biodyne FIREPol Master Mix (Tartu, Estonia, 04-12-00115). PCR products were purified and separated by electrophoresis on a 3% agarose gel and the amplified gene products were visualized under UV light after staining with ethidium bromide. Bands of 197 bp length were observed in all three replicates indicating successful amplification of the target gene fragment with no detected amplification in the no-template control confirming the absence of contamination or unspecific amplification. Moreover, Sanger sequencing of 2 PCR fragments was performed, resulting in 99.38% and 99.39% identity and 5e-73 and 4e-74 Error values respectively of the bla_*IMP-15*_ gene.

*Stenotrophomonas maltophilia* is a globally emerging multidrug-resistant pathogen. STM001 exhibits an extensively drug-resistant profile, due to its resistance against all CLSI-listed clinically relevant antibiotics except cefiderocol, which represents a last-resort option against such infections (13). It is evident that the recently adopted aztreonam + ceftazidime-avibactam combination disk testing shows synergy against this isolate as observed through disk diffusion testing, however, standard clinical breakpoints for such tests are lacking (21). Moreover, a recent 19-year retrospective study examining the antimicrobial-resistance patterns in *S. maltophilia* clinical isolates in Saudi Arabia reported an increase in the number of extensively-drug resistant isolates (22). Thus, establishing missing breakpoints for relevant antibiotic combinations and the development of novel treatments are required to face the increasing threat of elevated antimicrobial resistance rates of this bacterium.

On the other hand, the presence of *bla*_IMP-15_ in *S. maltophilia* is dangerous for two main reasons. First, ceftazidime and ceftazidime-avibactam are still considered potent treatment options for *Stenotrophomonas maltophilia* infections despite the presence of *bla*_L1_, which encodes intrinsic resistance to beta-lactam agents and carbapenems (23). However, the presence of the IMP-15 carbapenemase diminishes the efficacy of the drugs as they are primarily effective against non-metallo-beta-lactamase carbapenemases (24). Moreover, IMP-15 not only adds resistance to the strain itself, but it can also be transferred to other pathogens. The frequent occurrence of *S. maltophilia* with other multidrug-resistant pathogens in polymicrobial infections combined with its ability to transfer antimicrobial resistance genes to other bacteria (4, 6), offer favorable conditions for STM001 to transfer the carbapenemase IMP-15 to other pathogens. This horizontal gene transfer could compromise the effectiveness of carbapenems, thus promoting the spread of resistance in clinical settings.

To the best of our knowledge, this is the first time that *bla*_IMP-15_ was reported in *Stenotrophomonas maltophilia* worldwide. Studies on *Stenotrophomonas maltophilia* are relatively few compared to other pathogens. There have been very scarce reports on carbapenemases in *S. maltophilia* with most studies focusing on other resistance mechanisms. One study from Iran reported the presence of KPC, NDM, VIM, OXA-48 and IMP genes in *S. maltophilia* clinical isolates with IMP having the least frequent occurrence (4.39%) (25). This knowledge gap is particularly alarming since *S. maltophilia* is becoming of clinical concern worldwide, and the strains capable of transmitting carbapenem resistance to other pathogens present a major public health concern.

## References

1. Youenou B, Favre-Bonté S, Bodilis J, Brothier E, Dubost A, Muller D, et al. Comparative Genomics of Environmental and Clinical Stenotrophomonas maltophilia Strains with Different Antibiotic Resistance Profiles. Genome Biol Evol. 2015 Sep 1;7(9):2484–505.

2. Looney WJ, Narita M, Mühlemann K. Stenotrophomonas maltophilia: an emerging opportunist human pathogen. Lancet Infect Dis. 2009 May 1;9(5):312–23.

3. Brooke JS. Advances in the Microbiology of Stenotrophomonas maltophilia. Clin Microbiol Rev. 2021 May 26;34(3):10.1128/cmr.00030-19.

4. Brooke JS. Stenotrophomonas maltophilia: an Emerging Global Opportunistic Pathogen. Clin Microbiol Rev. 2012 Jan;25(1):2–41.

5. Stenotrophomonas maltophilia Susceptibility Testing Challenges and Strategies | Journal of Clinical Microbiology [Internet]. [cited 2025 Jan 20]. Available from: https://journals.asm.org/doi/full/10.1128/jcm.01094-21

6. Wang N, Tang C, Wang L. Risk Factors for Acquired Stenotrophomonas maltophilia Pneumonia in Intensive Care Unit: A Systematic Review and Meta-Analysis. Front Med [Internet]. 2022 Jan 12 [cited 2025 Jan 20];8. Available from: https://www.frontiersin.org/journals/medicine/articles/10.3389/fmed.2021.808391/full

7. Li Y, Liu X, Chen L, Shen X, Wang H, Guo R, et al. Comparative genomics analysis of Stenotrophomonas maltophilia strains from a community. Front Cell Infect Microbiol [Internet]. 2023 Nov 28 [cited 2025 Jan 16];13. Available from: https://www.frontiersin.org/journals/cellular-and-infection-microbiology/articles/10.3389/fcimb.2023.1266295/full

8. Xu Y, Cheng T, Rao Q, Zhang S, Ma YL. Comparative genomic analysis of Stenotrophomonas maltophilia unravels their genetic variations and versatility trait. J Appl Genet. 2023 May;64(2):351–60.

9. Yamagishi T, Matsui M, Sekizuka T, Ito H, Fukusumi M, Uehira T, et al. A prolonged multispecies outbreak of IMP-6 carbapenemase-producing Enterobacterales due to horizontal transmission of the IncN plasmid. Sci Rep. 2020 Mar 5;10(1):4139.

10. Weinstein MP, Lewis JS. The Clinical and Laboratory Standards Institute Subcommittee on Antimicrobial Susceptibility Testing: Background, Organization, Functions, and Processes. J Clin Microbiol. 2020 Feb 24;58(3):10.1128/jcm.01864-19.

11. EM100 Connect - CLSI M100 ED34:2024 [Internet]. [cited 2025 Jan 27]. Available from: https://em100.edaptivedocs.net/GetDoc.aspx?doc=CLSI%20M100%20ED34:2024&scope=user

12. Lin Q, Zou H, Chen X, Wu M, Ma D, Yu H, et al. Avibactam potentiated the activity of both ceftazidime and aztreonam against S. maltophilia clinical isolates in vitro. BMC Microbiol. 2021 Feb 22;21:60.

13. Kayama S, Kawakami S, Kondo K, Kitamura N, Yu L, Hayashi W, et al. In vitro activity of cefiderocol against carbapenemase-producing and meropenem-non-susceptible Gram-negative bacteria collected in the Japan Antimicrobial Resistant Bacterial Surveillance. J Glob Antimicrob Resist. 2024 Sep 1;38:12–20.

14. VM D, Peltzer A, Straub D, bot nf core, Wuennemann F, Garcia MU, et al. nfcore/bacass: v2.4.0 [Internet]. Zenodo; 2024 [cited 2025 Jan 21]. Available from: https://zenodo.org/records/14180424

15. Seemann T. Prokka: rapid prokaryotic genome annotation. Bioinforma Oxf Engl. 2014 Jul 15;30(14):2068–9.

16. Alcock BP, Huynh W, Chalil R, Smith KW, Raphenya AR, Wlodarski MA, et al. CARD 2023: expanded curation, support for machine learning, and resistome prediction at the Comprehensive Antibiotic Resistance Database. Nucleic Acids Res. 2023 Jan 6;51(D1):D690–9.

17. Feldgarden M, Brover V, Gonzalez-Escalona N, Frye JG, Haendiges J, Haft DH, et al. AMRFinderPlus and the Reference Gene Catalog facilitate examination of the genomic links among antimicrobial resistance, stress response, and virulence. Sci Rep. 2021 Jun 16;11:12728.

18. Crisan CV, Van Tyne D, Goldberg JB. The type VI secretion system of the emerging pathogen Stenotrophomonas maltophilia complex has antibacterial properties. mSphere. 8(6):e00584–23.

19. Multilocus Sequence Typing of Total-Genome-Sequenced Bacteria | Journal of Clinical Microbiology [Internet]. [cited 2025 Jan 24]. Available from: https://journals.asm.org/doi/10.1128/jcm.06094-11

20. Johansson MHK, Bortolaia V, Tansirichaiya S, Aarestrup FM, Roberts AP, Petersen TN. Detection of mobile genetic elements associated with antibiotic resistance in Salmonella enterica using a newly developed web tool: MobileElementFinder. J Antimicrob Chemother. 2021 Jan 1;76(1):101–9.

21. Mojica MF, Humphries R, Lipuma JJ, Mathers AJ, Rao GG, Shelburne SA, et al. Clinical challenges treating Stenotrophomonas maltophilia infections: an update. JAC-Antimicrob Resist. 2022 May 5;4(3):dlac040.

22. AlFonaisan MK, Mubaraki MA, Althawadi SI, Obeid DA, Al-Qahtani AA, Almaghrabi RS, et al. Temporal analysis of prevalence and antibiotic-resistance patterns in Stenotrophomonas maltophilia clinical isolates in a 19-year retrospective study. Sci Rep. 2024 Jun 24;14(1):14459.

23. Moriceau C, Eveillard M, Lemarié C, Chenouard R, Pailhoriès H, Kempf M. Stenotrophomonas maltophilia susceptibility to ceftazidime-avibactam combination versus ceftazidime alone. Med Mal Infect. 2020 May;50(3):305–7.

24. García-Castillo M, García-Fernández S, Gómez-Gil R, Pitart C, Oviaño M, Gracia-Ahufinger I, et al. Activity of ceftazidime-avibactam against carbapenemase-producing Enterobacteriaceae from urine specimens obtained during the infection-carbapenem resistance evaluation surveillance trial (iCREST) in Spain. Int J Antimicrob Agents. 2018 Mar 1;51(3):511–5.

25. Rizi KS, Jamehdar SA, Sasan MS, Ghazvini K, Aryan E, Safdari H, et al. Detection of extended-spectrum beta-lactamases and carbapenemases in clinical isolates of Stenotrophomonas maltophilia in the northeast of Iran. Gene Rep. 2024 Mar 1;34:101857.

